# iMab Antibody Binds Single-Stranded Cytosine-Rich Sequences and Unfolds DNA i-Motifs

**DOI:** 10.1101/2023.11.21.568054

**Authors:** Joseph Boissieras, Hugues Bonnet, Maria Fidelia Susanto, Dennis Gomez, Anton Granzhan, Eric Defrancq, Jérôme Dejeu

## Abstract

i-Motifs (iMs) are non-canonical, four-stranded secondary structures formed by stacking of hemi-protonated CH^+^·C base pairs in cytosine-rich DNA sequences, predominantly at pH < 7. The presence of iM structures in cells was a matter of debate until the recent development of iM-specific antibody, iMab, that was instrumental for several studies that suggested the existence of iMs in live cells and their putative biological roles. We assessed the interaction of iMab with cytosine-rich oligonucleotides by biolayer interferometry (BLI), pull-down assay and bulk-FRET experiments. Our results suggest that binding of iMab to DNA oligonucleotides is governed by the presence of runs of at least two consecutive cytosines and is generally increased in acidic conditions, irrespectively of the capacity of the sequence to adopt, or not, an iM structure. Moreover, the results of the bulk-FRET assay indicate that interaction with iMab results in unfolding of iM structures even in acidic conditions (pH 5.8 or 6.5), similarly to what has been observed with hnRNP K, well-studied single-stranded DNA binding protein. Taken together, our results suggest that iMab actually binds to blocks of 2–3 cytosines in single-stranded DNA, and call for more careful interpretation of results obtained with this antibody.

## INTRODUCTION

i-Motifs (iMs, or i-DNA) are non-canonical, four-stranded secondary structures formed by cytosine-rich DNA sequences upon mutual intercalation of hemi-protonated cytosine–cytosine base pairs (CH^+^·C), predominantly at relatively low pH (1–3). iM structures are known for three decades and well-characterized *in vitro* (4–9); nevertheless, the debate about the presence, persistence, and possible functions of these structures in cells is still relevant. It has been suggested that iMs play functional roles as transcription regulators (2, 10, 11) and therefore, could represent potential drug targets for a number of pathologies, including cancer and neurodegenerative diseases (11–14). The controversy regarding the presence of iMs in cells is mainly due to the strong pH dependence of their stability that, with a few exceptions (15, 16), may preclude their persistence at physiological pH and temperature. To explain this controversy, the existence of iM-stabilizing factors present in cells, such as molecular crowding or specific protein partners, has been postulated (17, 18). The molecular toolset to detect the presence of iMs in cells or interrogate their functions is sparse; thus, despite a massive research effort, no *bona fide* (i.e., truly specific and strongly stabilizing) small-molecule iM ligand has been identified to date (19, 20), which is in a stark contrast to a plethora of highly specific, biologically active G-quadruplex (G4) ligands. To address this gap, in 2018 the teams of D. Christ and M. Dinger raised an antibody (iMab) against the iM structure formed by the human telomeric C-rich strand (hTeloC) (21). The initial characterization of iMab demonstrated high affinity for hTeloC iM (*K*_D_ = 59 nM at pH 6.0, as *per* biolayer interferometry (BLI) analysis) and excellent selectivity, as no binding to the mutated sequence unable to form an iM structure (hTeloC-mut), double-stranded and hairpin DNA, or G4 structures was observed by using ELISA and BLI assays. At the same time, reduced but still non-negligible binding of iMab to the hTeloC strand was observed at pH 7.0 and even 8.0, which is rather surprising considering the transition pH of this sequence (pH_T_ = 6.11 at 20 °C) (22) and the fact that iM ↔ single-strand transitions are highly cooperative, meaning that no significant fraction of this sequence could be present in the iM form at pH 7.0 or, even less likely, at pH 8.0 (8). Despite this inconsistency, immunostaining of human cells with iMab clearly evidenced the accumulation of iMab foci in the nuclei, which was interpreted as the presence of iM structures in cells (21). This observation was subsequently extended to other species (23).

Since this seminal work, iMab was commercialized and rapidly found numerous applications as a molecular biology tool to study the formation and dynamics of iM structures in cells by immunofluorescence (23, 24), and to obtain genome-wide maps of iM-forming sequences. Along these lines, Christ and coll., using DNA immunoprecipitation and sequencing, identified in the human genome over 650, 000 sequences that were recognized by iMab and significantly enriched in cytosine; a biophysical validation of a subset of these sequences confirmed that most (but not all) were able to form iM structures *in vitro*, at pH 6.0 and 25 °C (25). In a more recent study, S. Richer and coll., using CUT&Tag technique, found that most iMab peaks (85– 94%) were located in promoter regions of genes and significantly co-localized with open-chromatin markers, which was consistent with the putative implication of iMs in transcription regulation (26). A biophysical validation confirmed that a number of sequences corresponding to iMab peaks were indeed able to form iM structures *in vitro* in acidic conditions (pH 5.4), but not in neutral (*i.e.,* close to physiological) conditions, thus keeping the conundrum regarding the factors that could potentially stabilize these structures in cells and allow their immunodetection with iMab.

In parallel, Schneekloth and coll. assessed the sequence preferences of iMab binding *in vitro* using a DNA microarray exposing over 11, 000 various DNA sequences, and found that binding of iMab was correlated with the length of C-tracts in DNA sequences (typically linked to higher stability of the corresponding iMs) (27). Interestingly, the correlation analysis performed in that study indicated high degree of similarity (Pearson’s *r* = 0.72) of the binding profile of iMab with that of hnRNP K, one of poly(C)- binding proteins (28, 29). The structural details of poly(C) binding by hnRNP K are known since at least two decades: this protein binds to the runs of at least three cytosines in single-stranded DNA regions, by making highly specific hydrogens bonds with unpaired cytosine residues (30–32). Considering the fact that within iM structures the cytosine residues participate in the formation of hemi-protonated CH^+^·C base pairs and become unavailable for additional hydrogen bonding, it is highly unlikely that hnRNP K could bind folded iMs. Accordingly, a recent single-molecule study by Wu and coll. provided solid evidence that hnRNP K actually unfolds iMs *in vitro*, namely through the binding to the corresponding single-strands (33).

Considering all these observations, we questioned the actual conformation of DNA sequences recognized by iMab antibody. To shed light on this riddle, we reassessed the interaction of hTeloC and its several variants, unable to adopt iM structures even in acidic conditions, with different forms of iMab (scFv-His_6_ and scFv-His_6_-FLAG) by using BLI and pull-down experiments. Our results provide firm evidence that binding of iMab to DNA oligonucleotides is governed by the presence of runs of at least two consecutive cytosines and is generally increased in acidic conditions, **irrespectively of the capacity of the sequence to adopt, or not, an iM structure**. Moreover, using bulk FRET experiments, we demonstrate that, similarly to hnRNP K, iMab is able to induce unfolding of iM structures in acidic conditions. These results strongly suggest that iMab actually binds single-stranded cytosine-rich DNA motifs.

## MATERIALS AND METHODS

The chemicals (Tris acetate and P20) were purchased from Sigma-Aldrich. The oligonucleotides were purchased from IDT or Eurogentec (HPLC purity grade). Two variants of the iMab antibody (scFv-His_6_, #Ab01462-30.11 and scFv-His_6_-FLAG, #Ab01462-30.135) were purchased from Absolute Antibody.

### BLI measurements

All BLI experiments were performed at 20 °C. BLI sensors coated with streptavidin (SA sensors) were purchased from Forte Bio (PALL). Prior to use, they were immerged for 10 min in a buffer (50 mM Tris-AcOH, pH 6 or 7.5, 0.05% v/v surfactant P20) to remove the protective sucrose layer from the sensor surface. The sensors were next dipped in a solution containing 100 nM of 3′-biotinylated DNA oligonucleotides during 900 s. The functionalized sensors were rinsed with the buffer for 10 min to remove the unbound molecules. The functionalized sensors were next dipped in the iMab solutions at different concentrations for 30 min interspersed by a rinsing step in the buffer solution for 10 min. Reference sensors without DNA immobilization were used to subtract the non-specific adsorption on the SA layer. One sensor was used per concentration due to the weak dissociation and to avoid using regenerative solution. The dissociation or association equilibrium constants, respectively *K*_D_ and *K*_A_, were calculated from the binding rate constants as *K*_D_ = *k*_off_/*k*_on_ or *K*_A_ = *k*_on_/*k*_off_ (where *k*_off_ and *k*_on_ represent the dissociation kinetic constant and the association kinetic constant, respectively). The reported values are the means of representative independent experiments and the errors provided are standard deviations from the mean. Each experiment was repeated at least two times.

### CD analysis

Circular dichroism studies were performed on a Jasco J-1500 spectropolarimeter using a 0.5 cm path length quartz cuvette. CD spectra were recorded at 20 °C using wavelengths range from 230 to 340 nm and were an average of four scans with a 0.5 s response time, a 1 nm data pitch, a 4 nm bandwidth and a 200 nm min_-1_ scanning speed. Samples containing 2.5 μM of DNA oligonucleotides were annealed at 90 °C in 10 mM LiAsMe_2_O_2_, 100 mM KCl buffer with indicated pH and cooled slowly overnight prior to CD analysis.

### Pull-down assay

5′-Biotinylated oligonucleotides were resuspended in 50 mM MES buffer (pH 5.8 or pH 7.0) at 1.5 µM final concentration, annealed at 95 °C for 5 min and cooled to room temperature overnight. Folded oligonucleotides were incubated with 100 ng of scFv-His_6_ iMab at 4 °C for 1 h in MES buffer with indicated pH (50 mM MES, 150 mM NaCl, 0.1% Triton and 1% BSA). After incubation, complexes were trapped with MES buffer-preequilibrated streptavidin-coated magnetic beads (Dynabeads™ M- 280 Streptavidin, ThermoFisher Scientific, #11205D). Excess antibody was washed by incubating 3 × 15 min with MES buffer without BSA, then magnetic beads were incubated with Laemmli buffer, heat-denatured at 95 °C for 10 min, and the dissociated fraction was recovered. Western blot was performed with anti-FLAG antibody 1:1000 (Sigma Aldrich, #F3165) incubated overnight at 4 °C in PBS-T buffer (PBS, 0.1% Tween 20) containing 1% BSA. Images were acquired using ChemiDoc imaging system (Bio-Rad).

### Bulk FRET experiments

The bulk FRET experiments were designed as described by Wu *et al.* (33). The sequences of oligonucleotides are given in **Supplementary Table S1**. A solution of Cy3- and Cy5-labelled strands (2 µM each) in a 50 mM phosphate (KH_2_PO_4_ / K_2_HPO_4_), 200 mM KCl, pH 5.8 buffer supplemented with 20% (w/v) PEG2000, in a total volume of 100 µL, was prepared and annealed for 5 min at 95 °C before being slowly brought to room temperature. Samples containing 4 nM of i-motif substrates in a total volume of 400 µL, accounting for the volume of the added iMab solution, were diluted in the same buffer adjusted to the desired pH. Samples were placed in Sigmacote-treated quartz cells with optical path lengths of 10 × 2 mm and thermostated at 20 °C. Fluorescence spectra were recorded with a HORIBA Jobin–Yvon FluoroMax-3 spectrofluorimeter, using excitation wavelength of 532 nm, emissions wavelength range of 550 nm to 800 nm (1-nm increment), excitation and emission bandwidths of 5 nm, and integration time of 0.2 s. After an initial measurement with DNA substrate alone, iMab was added to the desired final concentration, and fluorescence emission spectra were recorded in 2-min increments. FRET efficiency (*E*_FRET_) was calculated according to Equation (1):

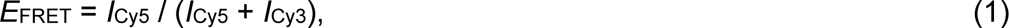

where *I*_Cy5_ and *I*_Cy3_ are the intensities of the Cy5 (667 nm) and Cy3 (565 nm) emission peaks, respectively.

## RESULTS

### BLI response of oligonucleotides is pH-dependent

In their seminal paper (21), Christ and coll. demonstrated, using BLI experiments, that iMab binds the iM-forming sequence hTeloC (*i.e.,* d[5′-TAA(CCCTTA)4-3′]) with an equilibrium constant (*K*_D_) close to 60 nM at pH 6.0. They suggested that iMab specifically recognizes the folded i-motif structure, because a gradual reduction of BLI signals was observed upon increasing pH above 6.0, interpreted as reduced binding of the antibody. However, using only the reduction of the BLI signal intensity can lead to misinterpretations. In fact, it is known that the amplitude of BLI signals strongly depends on the size and conformation of the interacting molecules (*i.e.* the one attached on the surface on the sensor as well as the analytes), because it is the induced change in optical thickness that gives rise to a BLI signal (34, 35). So, not only is molecular weight (MW) a determining factor, but also the molecular shape and packing density. In other words, if two molecules of the same MW show very different structures, they may well give rise to different BLI signals by virtue of their shape and packing ability. As an example, this phenomena has been used to study the unfolding of proteins (36). In the sensograms obtained by Christ and coll. at pH 6.0 (favorable for the formation of iMs) and at pH 8.0 (strongly unfavorable for iMs), the amplitude of the signal is different, but the shape is very similar for both the association and the dissociation steps. We anticipated that the difference of the signal amplitude could be not only due to the differences in binding of iMab, but rather due to a different conformation of the DNA sequence in these conditions.

To verify this point, we immobilized the biotinylated hTeloC sequence on BLI streptavidin (SA) sensors at pH 6.0 (conditions in which hTeloC is folded in a i-motif structure) and 7.5 (conditions in which hTeloC is unfolded at 20 °C). Remarkably, the intensity of the BLI response after the immobilization step was strongly pH-dependent (**Supplementary Figure S1**), which reflects a change in optical thickness. Stronger signals were obtained at low pH (6.0) which is compatible with the formation of the iM structure by hTeloC (cf. **Supplementary Figure S9B**). These results confirm that the amplitude of BLI signal is strongly influenced by the secondary structure of hTeloC oligonucleotides attached on the biosensor surface. **Consequently, the amplitude of BLI signals at a single concentration could not be used to correctly estimate the affinity, and a precise *K_D_* measurement using several concentrations of the analyte is required.** However, both the initial characterization of iMab (21) and the follow-up binding studies with various cytosine-rich sequences were performed with single analyte concentrations (25).

### iMab exhibits moderate selectivity between iM-folded and unfolded hTeloC sequence

We studied the binding of two variants of iMab (scFv-His_6_ and scFv-His_6_-FLAG) to hTeloC sequence at pH 6.0 and 7.5. A potentiometric titration of hTeloC (**Supplementary Figure S9A**) indicates that at pH 6.0 and 20 °C, 80% of this sequence exists in the iM form; conversely, at pH 7.5, the iM structure is fully unfolded, which is in agreement with the transition pH of this sequence (pH_T_ = 6.1) obtained by fitting of titration isotherms in this work and in other studies (22).

The raw sensograms obtained by dipping the sensor in solutions containing different concentrations of iMab are presented in **Figure 1** (scFv-His_6_) and **Supplementary Figure S2** (scFv-His_6_-FLAG). At pH 6.0, the amplitude of the BLI signal was similar to that observed in the Christ’s work (21), *i.e*., close to 10 nm in the presence of 500 nM iMab. For the same concentrations of iMab, the response was slightly lower at pH 7.5 (Δλ = 8 nm), but the shape of the curves was very similar. To calculate the equilibrium dissociation constants *K*_D_, a fitting by using a 1:1 model was applied in order to determine the kinetics constants (*k*_on_ and *k*_off_). A good agreement between the fitted and the experimental curves was observed.

**Figure 1.**
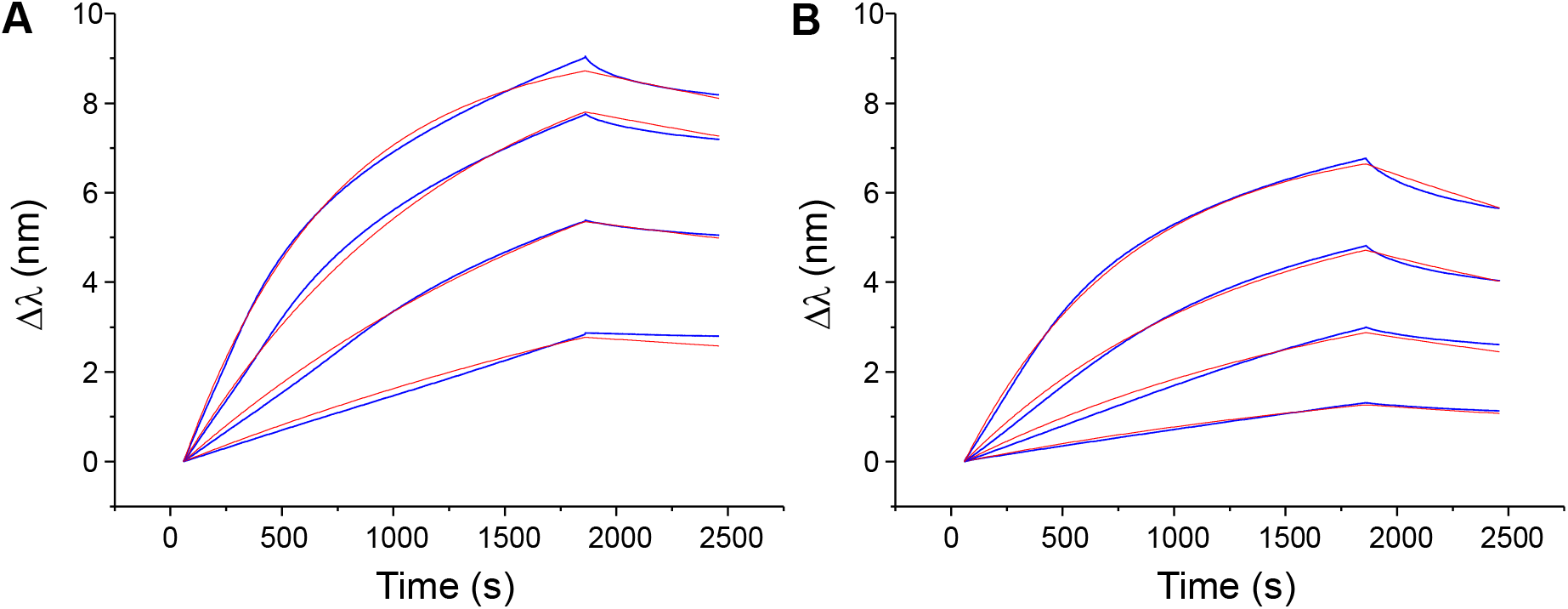
BLI analysis of binding of different concentrations of iMab scFv-His6 (62.5, 125, 250 and 500 nM) to hTeloC at pH 6.0 (**A**) and pH 7.5 (**B**). Blue lines are experimental curves; red lines are the fits to the 1:1 model.

The kinetic and thermodynamic parameters obtained for the two variants of iMab (scFv-His_6_ and scFv-His_6_-FLAG) at pH 6.0 and 7.5 are summarized in **Table 1**. Interestingly, for scFv-His_6_ the kinetic constants and the *K*_D_ values at pH 6.0 and 7.5 are essentially of the same order of magnitude (*K*_D_ = 34 and 108 nM, respectively). The resulting apparent selectivity (∼3.2-fold) is mainly due to a lower kinetic dissociation constant at pH 6.0. scFv-His_6_-FLAG has a slightly better affinity at pH 6.0 (*K*_D_ = 20 nM) and better apparent selectivity (∼8.2-fold) with respect to pH 7.5. However, in both cases, **the binding is non-negligible at pH 7.5, despite the fact that in the absence of iMab, iM structures formed by hTeloC sequence are fully unfolded in these conditions**.

**Table 1.**
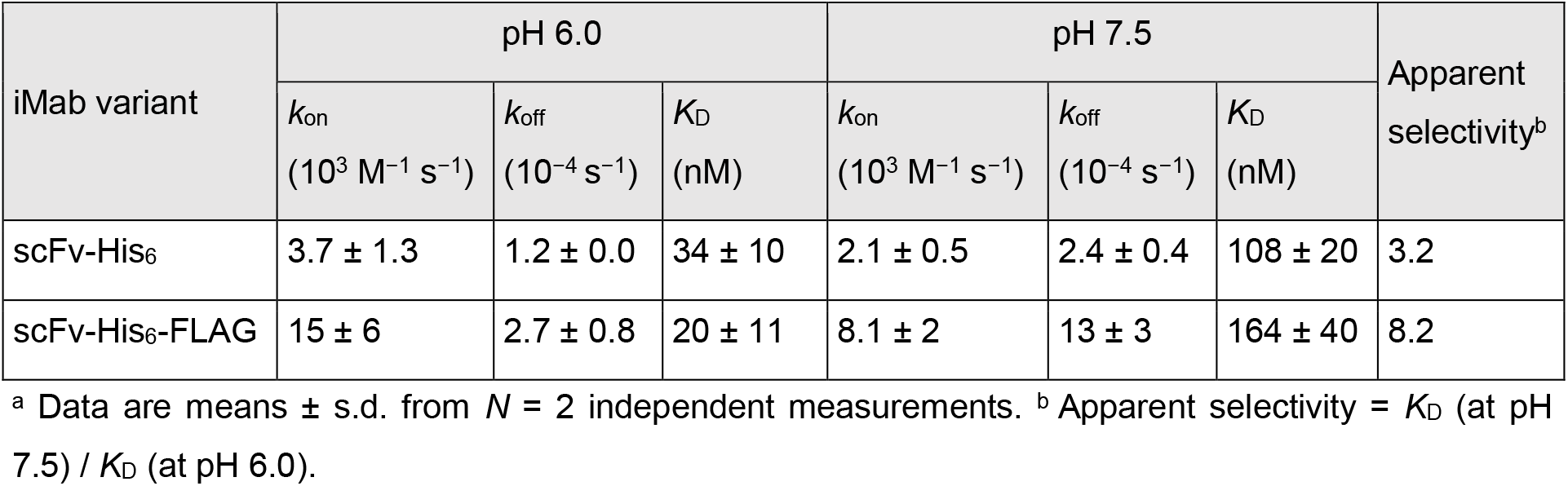
Kinetics constants (*k*_on_ and *k*_off_), calculated dissociation equilibrium constants (*K*_D_) and apparent selectivity for two variants of iMab antibody interacting with hTeloC sequence at pH 6.0 and 7.5._a_.

| iMab variant | pH 6.0 |  |  | pH 7.5 |  |  | Apparent selectivity<sup>b</sup> |
| --- | --- | --- | --- | --- | --- | --- | --- |
| | $k_{on}$<br>($10^3 \text{ M}^{-1} \text{ s}^{-1}$) | $k_{off}$<br>($10^{-4} \text{ s}^{-1}$) | $K_D$<br>(nM) | $k_{on}$<br>($10^3 \text{ M}^{-1} \text{ s}^{-1}$) | $k_{off}$<br>($10^{-4} \text{ s}^{-1}$) | $K_D$<br>(nM) | |
| scFv-His<sub>6</sub> | $3.7 \pm 1.3$ | $1.2 \pm 0.0$ | $34 \pm 10$ | $2.1 \pm 0.5$ | $2.4 \pm 0.4$ | $108 \pm 20$ | 3.2 |
| scFv-His<sub>6</sub>-FLAG | $15 \pm 6$ | $2.7 \pm 0.8$ | $20 \pm 11$ | $8.1 \pm 2$ | $13 \pm 3$ | $164 \pm 40$ | 8.2 |

<sup>a</sup> Data are means ± s.d. from $N = 2$ independent measurements. <sup>b</sup> Apparent selectivity = $K_D$ (at pH 7.5) / $K_D$ (at pH 6.0).

### iMab shows no preferential binding to the constrained iM scaffold

Previously, we described a constrained analogue (**1**) of hTeloC iM, assembled on a rigid cyclopeptide scaffold (**Figure 2A**). The particularity of this constrained iM is its higher stability; thus, (**1**) is folded at pH 6.5, while the native sequence is unfolded by ∼90% in these conditions (**Figure 2B** and **Supplementary Figure S9B**) (19, 37). The interaction of the constrained hTeloC (**1**) immobilized on SA sensor with iMab (scFv-His_6_-FLAG) was studied at pH 6.5 in comparison with the native sequence; despite a weaker signal, the recognition pattern for the iMab was similar to the previous experiments (**Supplementary Figure S3**). After the fitting of the sensograms, the kinetic constants were determined and the *K*_D_ value was calculated (**Table 2**). Remarkably, iMab binds to the constrained hTeloC (**1**), which *a fortiori* adopts an iM structure at pH 6.5, with a 3-fold *lower* affinity comparing with the native sequence, which is chiefly unfolded in these conditions (*K*_D_ = 155 and 54 nM, respectively). This suggests that binding of iMab is not driven by the iM structure.

**Figure 2.**
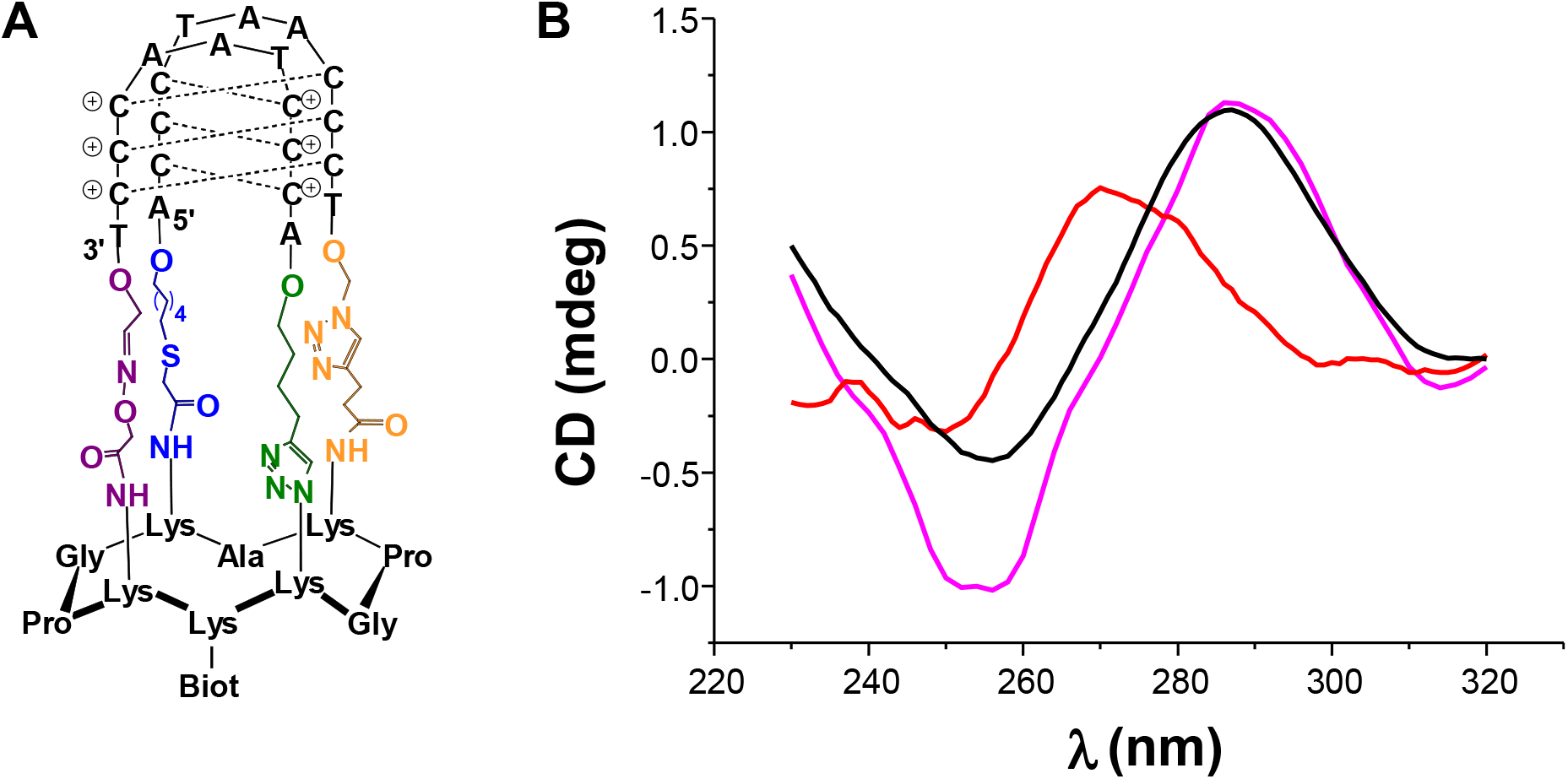
(**A**) Structure of the constrained iM (**1**). (**B**) CD spectra recorded at 20 °C for constrained (black) and native hTeloC (red) substrates at pH 6.5, and native hTeloC at pH 6.0 (magenta).

**Table 2.** Kinetic constants (*k*_on_ and *k*_off_) and calculated dissociation equilibrium constants (*K*_D_) for interaction of iMab (scFv-His_6_-FLAG) with the native and the constrained HTeloC substrates at pH 6.5 and 20 °C._a_.

| Substrate | $k_{on}$ ($10^3 \text{ M}^{-1} \text{ s}^{-1}$) | $k_{off}$ ($10^{-4} \text{ s}^{-1}$) | $K_D$ (nM) |
| --- | --- | --- | --- |
| Native hTeloC | $11 \pm 0.3$ | $6.1 \pm 0.9$ | $54 \pm 6$ |
| Constrained hTeloC (1) | $4.6 \pm 0.3$ | $7.2 \pm 0.8$ | $155 \pm 22$ |

<sup>a</sup> Data are means ± s.d. from $N = 2$ independent measurements.

### iMab binds cytosine-rich sequences unable to adopt iM structures

We used BLI to assess the binding of iMab to a panel of 14 DNA oligonucleotides (**Table 3**). In this panel, eleven sequences have the same length as hTeloC (27 nt) but differ by the number and the arrangement of cytosine residues critical for the formation of the iM structure; in particular, three variants (hTeloC-scr, hTeloC-3x4 and hTeloC-2x6T) have the same number of cytosine residues (*i.e.,* 12) as the native sequence. Additionally, we included the two sequences used by Christ and coll. as negative controls (hTeloC-X3 and hTeloC-mut) (21), as well as one hairpin (hp-ATT) and a cytosine-poor single-stranded sequence (ss-DNA). A close inspection of the corresponding CD spectra revealed that among these sequences, two variants, namely hTeloC-3x4 and hTeloC-2x6T, were able to adopt iM structures similarly to hTeloC, as indicated by the iM-characteristic CD bands at 283–285 nm observed at pH 6.0 and 5.5 (**Supplementary Figure S10**). In contrast, 11 other sequences were unable to form iM structures, as their CD spectra remained essentially identical upon decreasing pH from 7.5 to 6.0 and even to 5.5, except for hTeloC-3x3 and hTeloC-X3 (both having very similar sequence) whose CD spectra were minimally affected at pH 5.5 (but not at pH 6.0), allowing to suggest the presence of minor fractions of iM structures in these conditions.

**Table 3.**
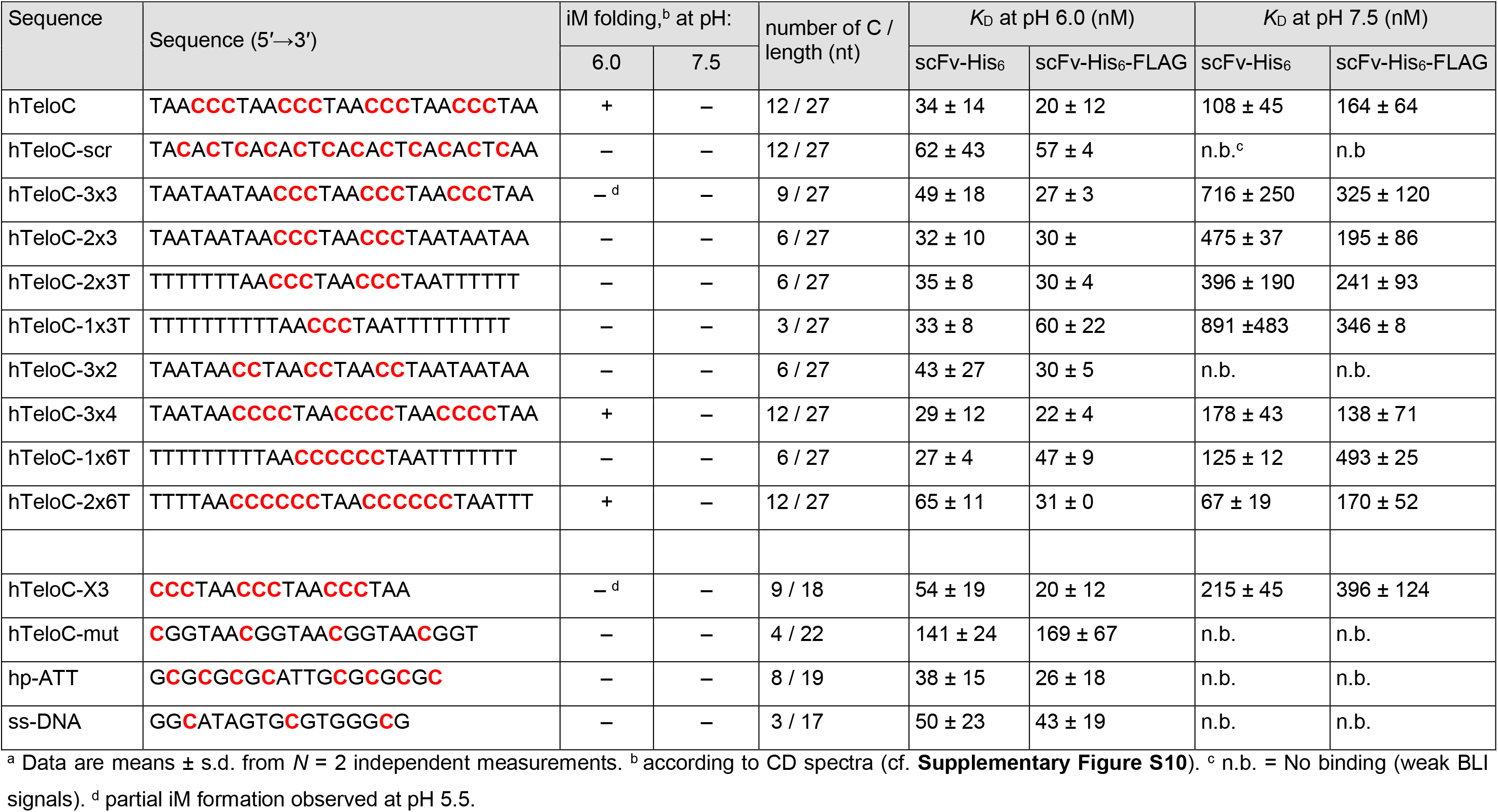
Modified sequences used in this study and the corresponding *K*_D_ values for the two variants of iMab, obtained from BLI experiments at pH 6.0 or 7.5 and 20 °C._a_.

The *K*_D_ values obtained from BLI experiments with two variants of iMab (scFv-His_6_ and scFv-His_6_-FLAG) are summarized in **Table 3** (sensograms and 1:1 model fits, **Supplementary Figures S4–S8**). The *K*_A_ (= 1 / *K_D_*) values for scFv-His_6_ and scFv-His_6_-FLAG are plotted on **Figure 3** to facilitate the comparison. Remarkably, at pH 6.0, the *K*_D_ values obtained for most sequences were essentially in the same range (for scFv-His_6_, 27–65 nM; for scFv-His_6_-FLAG, 20–60 nM) with an exception for hTeloC- mut for which significantly higher *K*_D_ values were observed (141 and 169 nM for scFv-His_6_ and scFv-His_6_-FLAG, respectively). Conversely, at pH 7.5, the affinity was strongly reduced for *all* sequences, with no binding (*i.e.,* weak BLI response) observed for hp-ATT, ss-DNA, HteloC-mut, HTeloC-scr, and HTeloC-3x2, that is, **all sequences that do not contain at least two consecutive cytosines**. Most strikingly, in both pH conditions, these data suggest no significant difference, in terms of binding affinity, between the sequences capable of forming iM structures (hTeloC, hTeloC-3x4, and hTeloC-2x6T) and those unable of iM folding even in most favorable conditions. Instead, they indicate that **i) iMab preferentially binds DNA oligonucleotides containing at least two or, better, three consecutive cytosines**, and **ii) for all oligonucleotides (whether they are able to adopt an iM structure or not), the binding is strongly increased in acidic conditions**.

**Figure 3.**
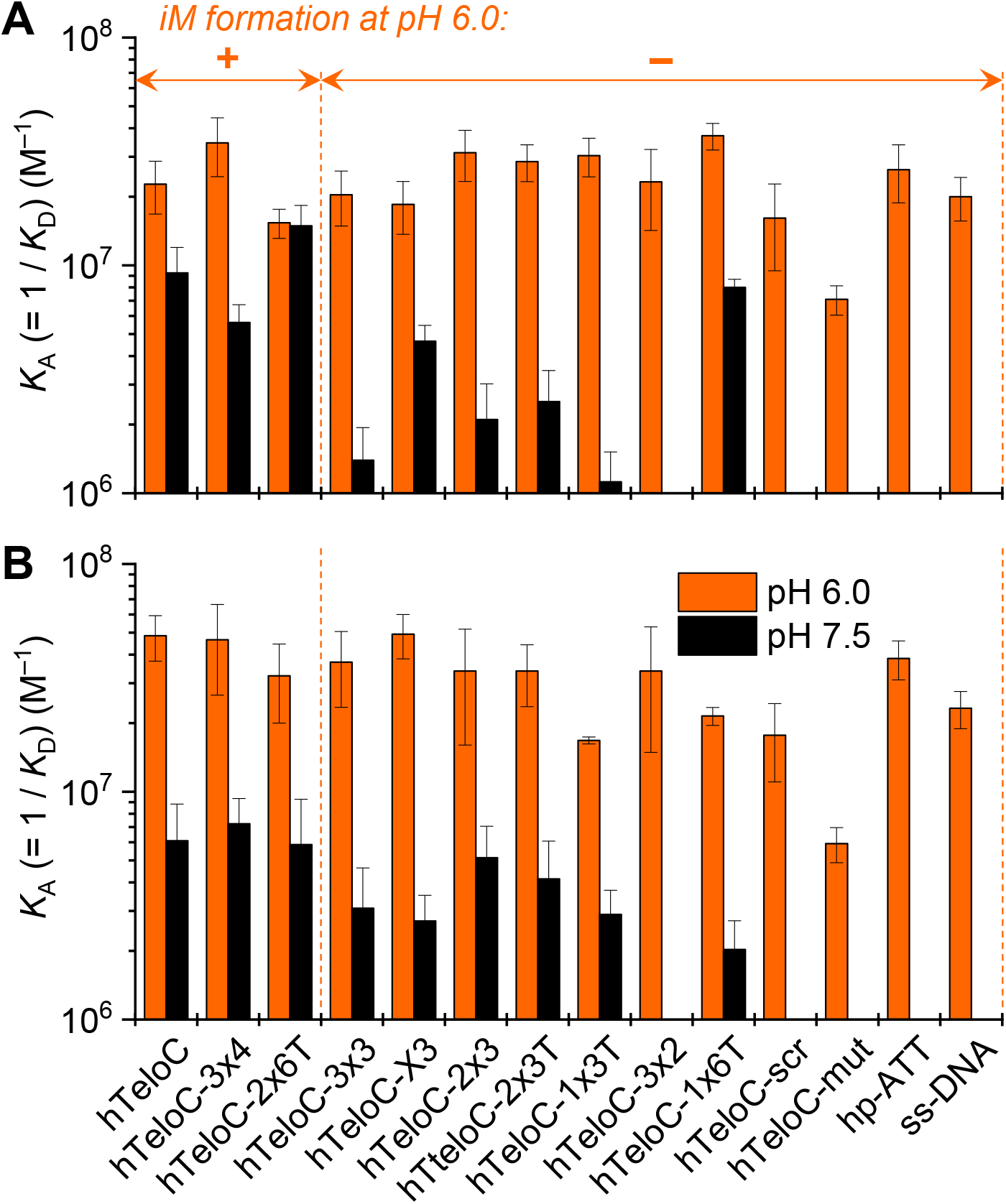
Comparison of *K*_A_ (= 1/*K*_D_) values for binding of iMab scFv-His_6_ (**A**) and scFv-His_6_-FLAG (**B**) to the oligonucleotides listed in Table 3 at pH 6.0 (orange) and 7.5 (black bars). Data are means ± s.d. from two independent experiments. The capacity of oligonucleotides to adopt iM structures at pH 6.0 is inferred from CD data.

Additionally, the interaction of iMab (scFv-His_6_-FLAG) with four variants of the telomeric sequence (hTeloC, hTeloC-3x3, hTeloC-3x4 and hTeloC-3x2) was studied through a pull-down assay followed by the detection of oligonucleotide-bound and subsequently released antibody by Western blotting. The results demonstrated strong binding of scFv-His_6_-FLAG iMab to all four sequence variants in acidic conditions (pH 5.8, **Figure 4A**), independently of their ability to form iM structures in these conditions (cf. **Table 3**). Remarkably, the interaction of iMab with all telomeric variants was also observed at pH 7.0, where none of the sequences could adopt an iM structure (**Figure 4B**). Of note, in acidic conditions iMab showed slightly weaker binding to hTeloC-3x2 compared to the native hTeloC sequence, as indicated by the intensities of the corresponding bands; however, similar differences were also observed at neutral pH, once again suggesting that iM structure is not the driver of iMab binding.

**Figure 4.**
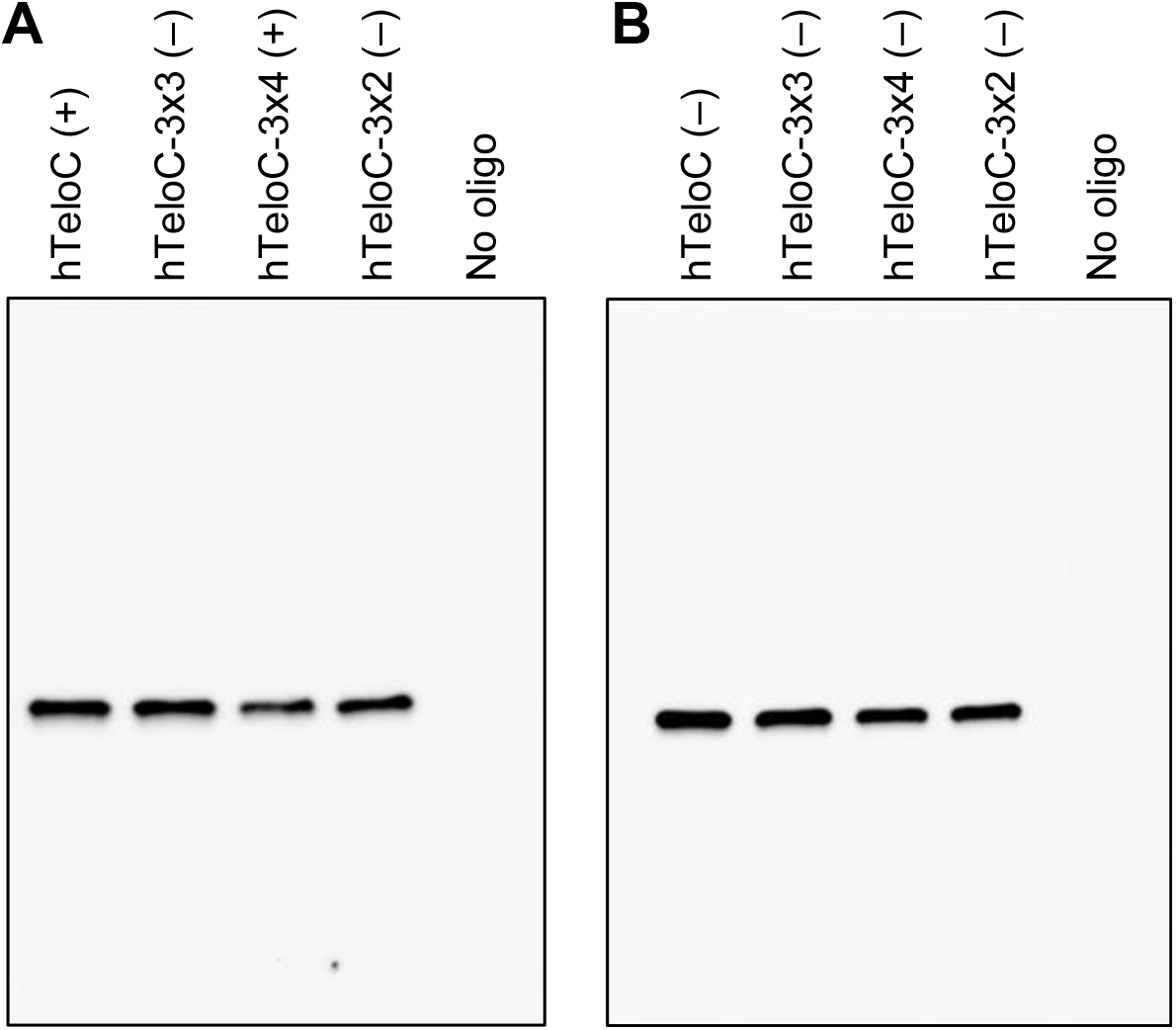
Pull-down following by Western blot detection of iMab scFv-His_6_-FLAG, performed with 5ʹ-biotinylated oligonucleotides at pH 5.8 (**A**) and at pH 7.0 (**B**). (+) and (–) indicate the capacity of the oligonucleotide to adopt iM structure in the experimental conditions.

#### iMab unfolds iM structures in bulk-FRET experiments

Wu and coll. recently demonstrated, using single-molecule and bulk-FRET experiments, that hnRNP K, a poly(C)-binding protein (30–32, 38), rapidly unfolds an iM structure through a stepwise process driven by the binding of protein to the cytosine-rich single strand of DNA (33). We employed the bulk-FRET assay to assess the capacity of iMab to induce unfolding of iMs. In this assay, the iM-forming sequence is 5′-labeled with a donor fluorophore (Cy3) and hybridized, *via* a 17-nt 3′-overhang, to a complementary strand labelled with an acceptor fluorophore (Cy5) in the vicinity of the iM–duplex junction (**Figure 5A**). In the folded state, the proximity between the fluorophores results in high efficiency of FRET (*E*_FRET_); conversely, unfolding of the iM upon binding of the protein leads to a larger distance between the fluorophores and a decrease of *E*_FRET_ value. Using the same iM-forming sequence as Wu and coll. (Py25, derived from the *MYC* promoter, **Table S1**), we first assessed the effect of the recombinant hnRNP K (800 nM) at pH 5.8 and observed a rapid increase of Cy3 fluorescence with a concomitant decrease of Cy5 fluorescence, consistent with the reduction of *E*_FRET_ from 0.74 to 0.36 after 3 min (**Figure S11**), in agreement with the literature data (33). Importantly, the same effect was observed upon addition of iMab. Thus, addition of 400 nM iMab scFv-His_6_ to Py25 iM substrate resulted in a decrease of *E*_FRET_ from 0.78 to 0.55 after 100 min. Addition of 800 nM iMab led to an even more rapid and stronger decrease of *E*_FRET_ (to 0.35), demonstrating that this effect is concentration-dependent (**Figure 5B**). Remarkably, the kinetics of the iMab-driven unfolding was dramatically accelerated at pH 6.5 (**Figure 5C**). Similar unfolding, albeit with a slower rate, was observed with the hTeloC sequence at pH 5.8 (**Figure 5D**). Finally, antibody G4-specific BG4 did not unfold Py25 iM (**Figure 5E**), giving evidence that the unfolding process is specific for the iMab antibody that behaves, in this regard, similarly to hnRNP K protein.

**Figure 5.**
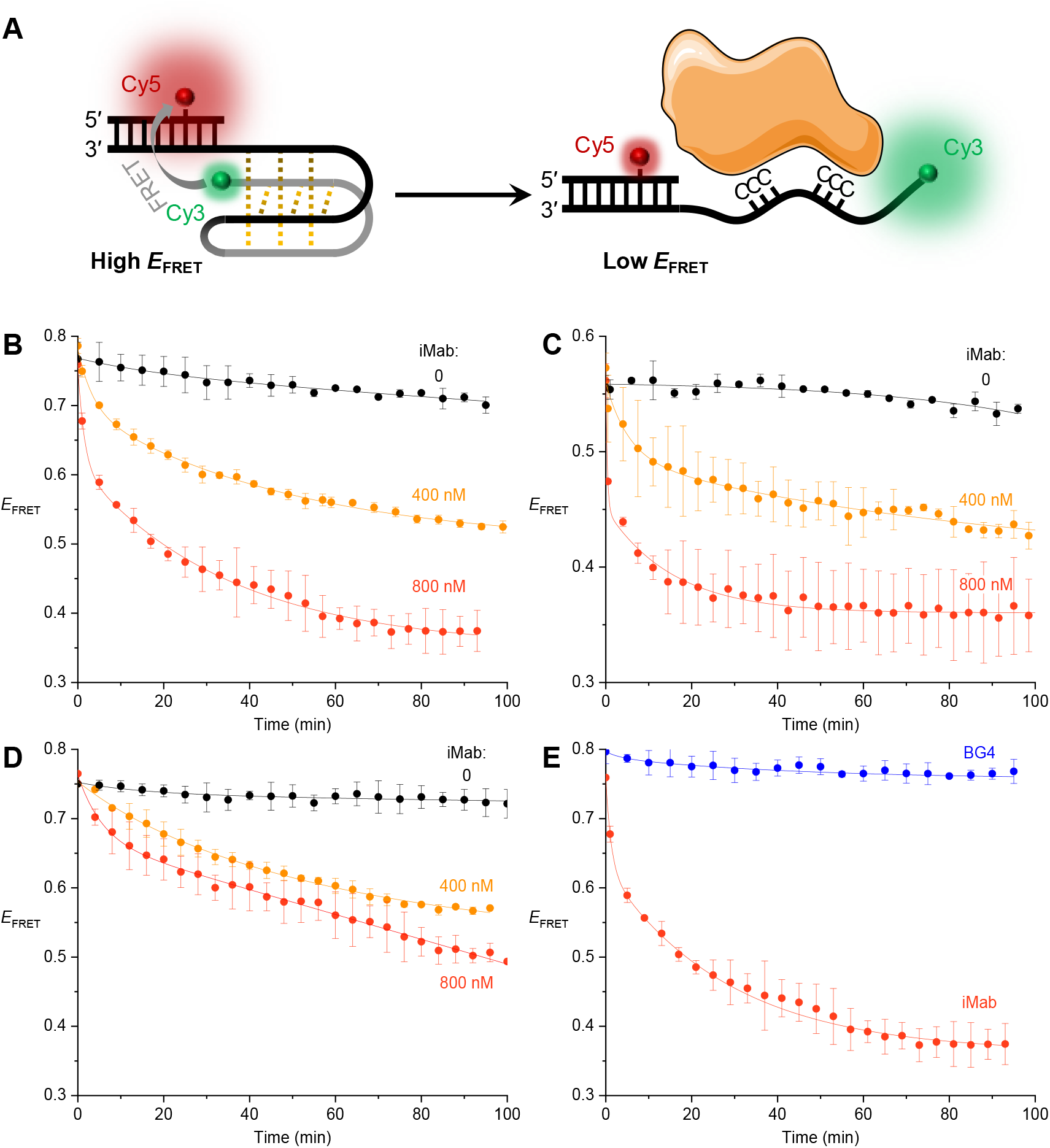
(**A**) Principle of the bulk-FRET assay to assess the unfolding of iM structures by proteins. (**B**–**C**) Time-dependent change of *E*_FRET_ upon addition of iMab scFv-His_6_ (0, 400, or 800 nM) to Py25 substrate at pH 5.8 (**B**) or pH 6.5 (**C**). (**D**) Time-dependent change of *E*_FRET_ upon addition of iMab scFv-His_6_ (0, 400 or 800 nM) to hTeloC substrate at pH 5.8. (**E**) Comparison of effects of BG4 and iMab antibodies (800 nM each) on the unfolding of Py25 substrate at pH 5.8. In all experiments, the concentration of the iM substrate was 4 nM in 50 mM phosphate buffer containing 200 mM KCl and 20% (w/v) PEG2000.

## DISCUSSION

In this work, using BLI analysis, we systematically investigated the binding of two commercially available variants of iMab antibody (scFv-His_6_ and scFv-His_6_-FLAG) to cytosine-rich DNA oligonucleotides. Initially, in the experiments with the native hTeloC sequence (*i.e.,* the sequence against which iMab was originally raised), we observed limited selectivity for binding at pH 6.0 (where this sequence predominantly adopts an iM structure) comparing to pH 7.5 where this sequence is fully unfolded. This moderate selectivity (3.2-fold and 8.2-fold for scFv-His_6_ and scFv-His_6_-FLAG, respectively), already observed in the original study, is rather surprising considering the extremely weak binding of iMab to all other DNA structures (G4s, DNA hairpins and single-stranded RNA) (21). Two alternative scenarios could explain this moderate selectivity: (i) iMab binds and strongly stabilizes iM structures to the extent that they persist as such in the conditions of BLI experiments at pH 7.5 and 20 °C; (ii) iMab binds to the unfolded hTeloC sequence and thereby unfolds iM structures that are otherwise stable at pH 6.0. Further sets of experiments allowed us to discriminate between these two possibilities.

First, we assessed the binding of iMab to the constrained iM structure (**1**) assembled on a cyclopeptide scaffold. Being significantly more stable than the native iM, this structure is very similar to the iM formed by the native hTeloC sequence, featuring the same iM core consisting of six CH_+_–C base pairs and two AAT loops. To our surprise, the binding affinity of iMab to this structure was 3-fold lower compared to the native hTeloC sequence (which is mainly unfolded in the conditions of this experiment, i.e., at pH 6.5), suggesting that **folded iM structure is not the driver of iMab binding**. Of note, one of the sides of (**1**) is hindered by the surface-linked cyclopeptide, which may prevent antibody binding. However, in BLI experiments the biotinylated, surface-bound 3′-end of the native hTelo iM is similarly unavailable for antibody binding, a fact which minimizes the differences between these two substrates.

In line with these observations, additional BLI experiments did not reveal any significant difference in iMab binding between the sequences capable of forming an iM structure (hTeloC, hTeloC-3x4 and hTeloC-2x6T) and 10 other DNA sequences of similar length, but unable to form iMs even in most favorable conditions (*i.e.,* at pH 5.5). The only sequence that demonstrated a significantly reduced iMab binding was hTeloC-mut, which was previously used as a negative control by Christ and coll. However, the cytosine residues in this sequence are few (4 *vs*. 12 in hTeloC) and spatially isolated. Remarkably, other sequence variants with similar number of cytosines, but assembled in blocks of two or three (hTeloC-1x3T, hTeloC-3x2) demonstrated *K*_D_ values close to that of hTeloC at pH 6.0. In all cases, the binding of iMab was strongly reduced at pH 7.5, suggesting that it primarily depends on **(i) pH (stronger binding in acidic conditions, *independently* of the capacity of the sequence to form iM)** and **(ii) the presence of *blocks* of at least two or, better, three cytosines**. Furthermore, the results of the pull-down assay performed in two pH conditions also demonstrated the incapacity of iMab to discriminate between the sequences capable (hTeloC, hTeloC-3x4) or not (hTeloC-3x3, hTeloC-3x2) of folding into iM structures, further consolidating the above conclusions.

Finally, the results of bulk-FRET assay clearly indicate that iMab binds and unfolds iM structures even in acidic conditions, where iMs are otherwise thermodynamically stable. This implies that the energy of iMab interaction with unfolded cytosine-rich sequences is superior to the thermodynamic stability of iMs. Altogether, all these observations can be rationalized by a model where iMab binds to blocks of cytosines in unfolded DNA sequences. This interaction may be similar to the binding mode of KH3 domains of hnRNP K to cytosine-rich DNA strands, but likely involves amino acid residues whose p*K*_a_ is closer to the physiological conditions (pH 6–8), instead of two arginines in the KH3 domain of hnRNP K (31). In view of this model, addition of iMab at pH 6.0 leads to unfolding of iMs both in bulk-FRET and BLI experiments; however, the binding is reduced at increased pH, presumably due to the deprotonation of the key amino acid residues in iMab, resulting in the “apparent” selectivity as observed for hTeloC sequence.

Last but not least, our model in no way contradicts the experimental results observed in previous studies employing iMab, but rather calls for a different interpretation of such data. Thus, the microarray study of Schneekloth and coll. included > 10, 000 iM-forming sequences containing blocks of 2 to 10 cytosines, but only 500 “negative controls” where C-tracts were replaced by adenines or thymines; it is thus unsurprising that the authors evidenced strong binding of iMab to cytosine-rich, and no binding to cytosine- poor sequences, whatever pH condition used (27). Likewise, the iMab peaks identified in the CUT&Tag analysis by Richter and coll. were subject to a bioinformatics filter through a regular expression comprising four to five blocks of at least two cytosines. Expectedly, a number of recovered sequences were shown to fold iM structures *in vitro* (26). Interestingly, the high number of iMab peaks observed by Richer and coll. in physiological conditions and the fact that iMab peaks are amplified in close proximity of BG4 peaks, is biologically plausible if one accepts that iMab binds single-stranded cytosine-rich regions, that become available for the antibody binding, for example upon formation of G-quadruplexes on the opposite DNA strand in open chromatin. Likewise, the iMab foci observed in immunostaining experiments should be rather interpreted as an evidence of single-stranded, cytosine-rich DNA regions. In line with this hypothesis, a recent in-cell NMR study by Trantirek and coll. demonstrated that iM-forming sequences with pH_T_ < 7 are predominantly unfolded in cells, suggesting that only a minor fraction of iM-forming sequences could form iM structures in physiological conditions (39).

## ACKNOWLEDGEMENTS

The authors thank Dr. Jean-Louis Mergny (*École Polytechnique*, Palaiseau) and Prof. Jean-François Riou (MNHN, Paris) for stimulating discussions and critical reading of the manuscript. The NanoBio-ICMG platforms (UAR 2607) are acknowledged for their support.

