## Supplementary Information for "iMab Antibody Binds Single-Stranded Cytosine-Rich Sequences and Unfolds DNA i-Motifs"

#### Table of contents

**BioLayer Interferometry (BLI) studies**

**Figure S1.** BLI response during hTeloC immobilization at pH 6.0 and 7.5 S2

**Figure S2.** Sensorgrams obtained for interaction of iMab scFv-His$_6$-FLAG with hTeloC at pH 6.0 and pH 7.5 S2

**Figure S3.** Sensorgrams obtained for interaction of iMab with native hTeloC and the constrained system **(1)** S2

**Figures S4–S7.** Sensorgrams obtained for interaction of iMab with various variants of hTeloC sequence S3–S6

**Figure S8.** Sensorgrams obtained for interaction of iMab with hp-ATT and ss-DNA. S7

**CD studies**

**Figure S9.** pH Titration and CD spectra of native hTeloC at pH 6.0, 6.5 and pH 7.5 S8

**Figure S10.** CD spectra of various variants of hTeloC sequence S9

**Bulk-FRET experiments**

**Table S1.** Oligonucleotides used in bulk-FRET experiments S10

**Figure S11.** Time-dependent variation of fluorescence emission spectra. S10

### BLI studies

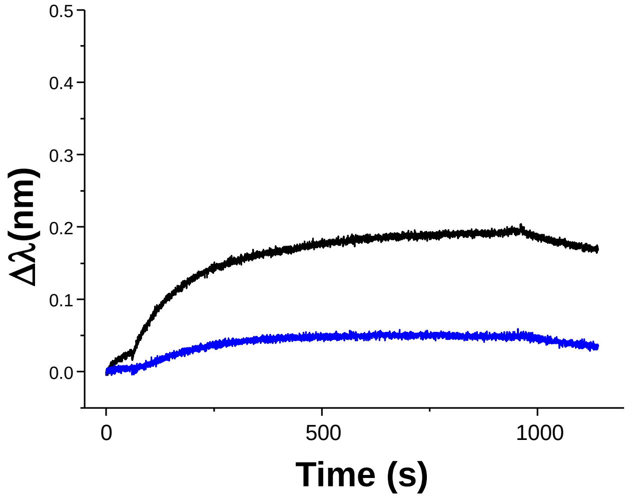


**Figure S1.** BLI signals observed during immobilization of native hTeloC at pH 6.0 (black) and 7.5 (blue).

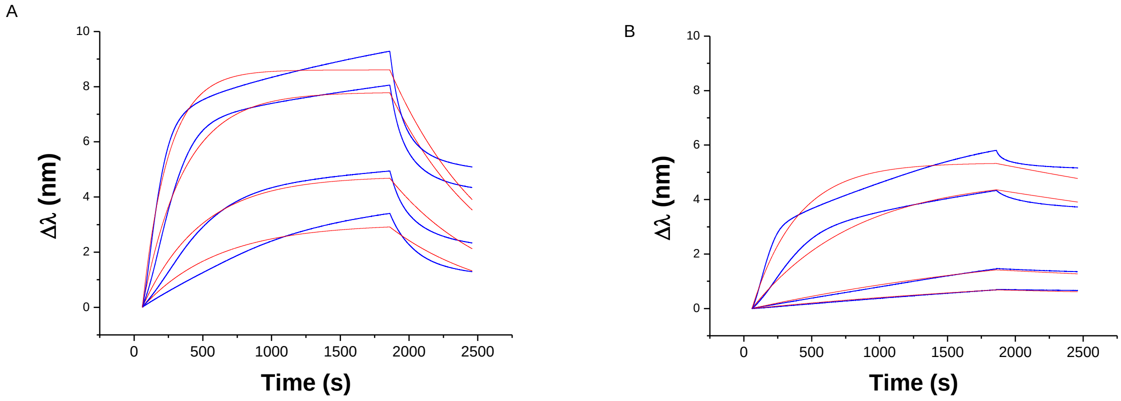


**Figure S2.** BLI analysis of binding of different concentrations of iMab scFv-His$_6$-FLAG (62.5, 125, 250 and 500 nM) to hTeloC at pH 6.0 (**A**) and pH 7.5 (**B**). Blue lines are experimental curves; red lines are the fits to the 1:1 model.

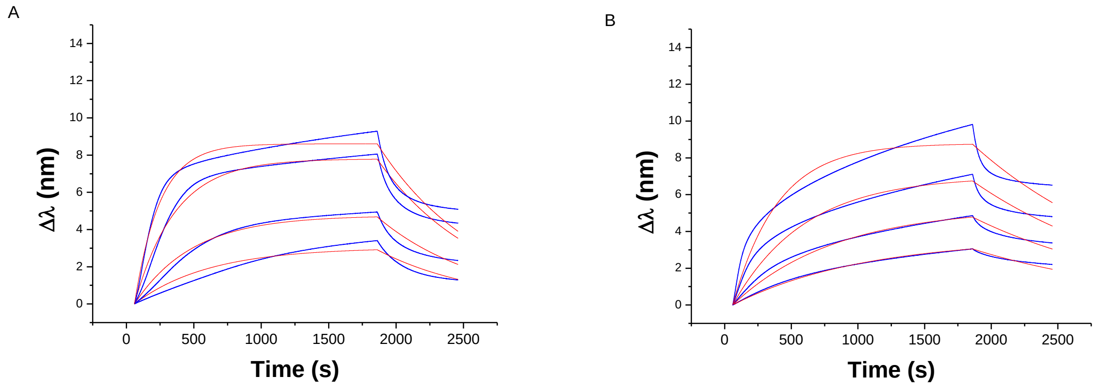


**Figure S3**: BLI analysis of binding of different concentrations of iMab scFv-His6-FLAG (62.5, 125, 250 and 500 nM) to native hTeloC (**A**) and constrained hTeloC **1** (**B**) at pH 6.5. Blue lines are experimental curves; red lines are the fits to the 1:1 model.

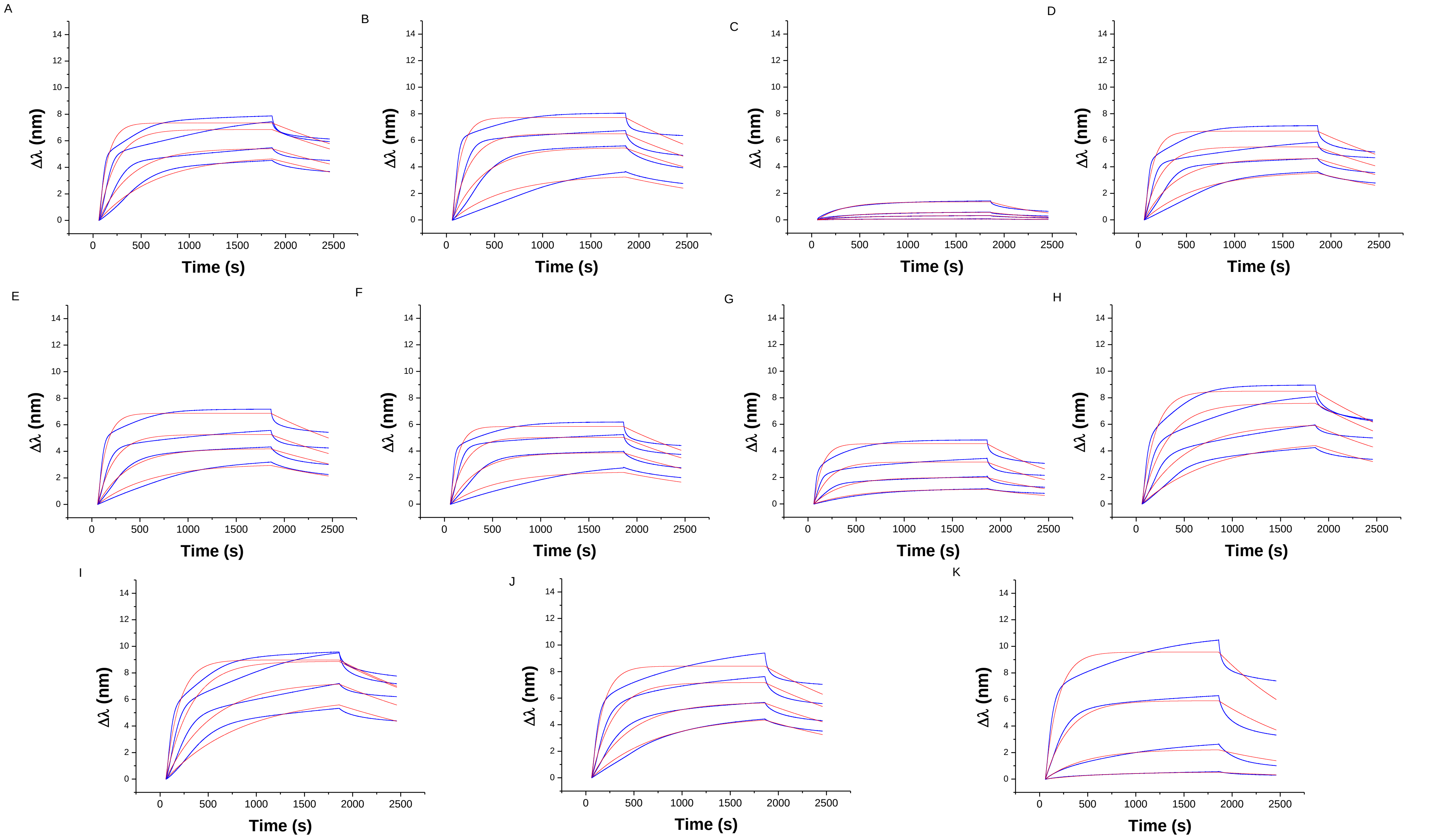


**Figure S4.** BLI analysis of binding of different concentrations of iMab scFv-His$_6$-FLAG (62.5, 125, 250 and 500 nM) to hTeloC (**A**), hTeloC 3x3 (**B**), hTeloC X3 (**C**), hTeloC 2x3 (**D**), hTeloC 3x2 (**E**), hTeloC 2x3T (**F**), hTeloC 1x3T (**G**), hTeloC 1x6 T (**H**), hTeloC 2x6T (**I**), hTeloC-Mut (**J**), and hTeloC-scr (**K**) at pH 6.0. Blue lines are experimental curves; red lines are the fits to the 1:1 model.

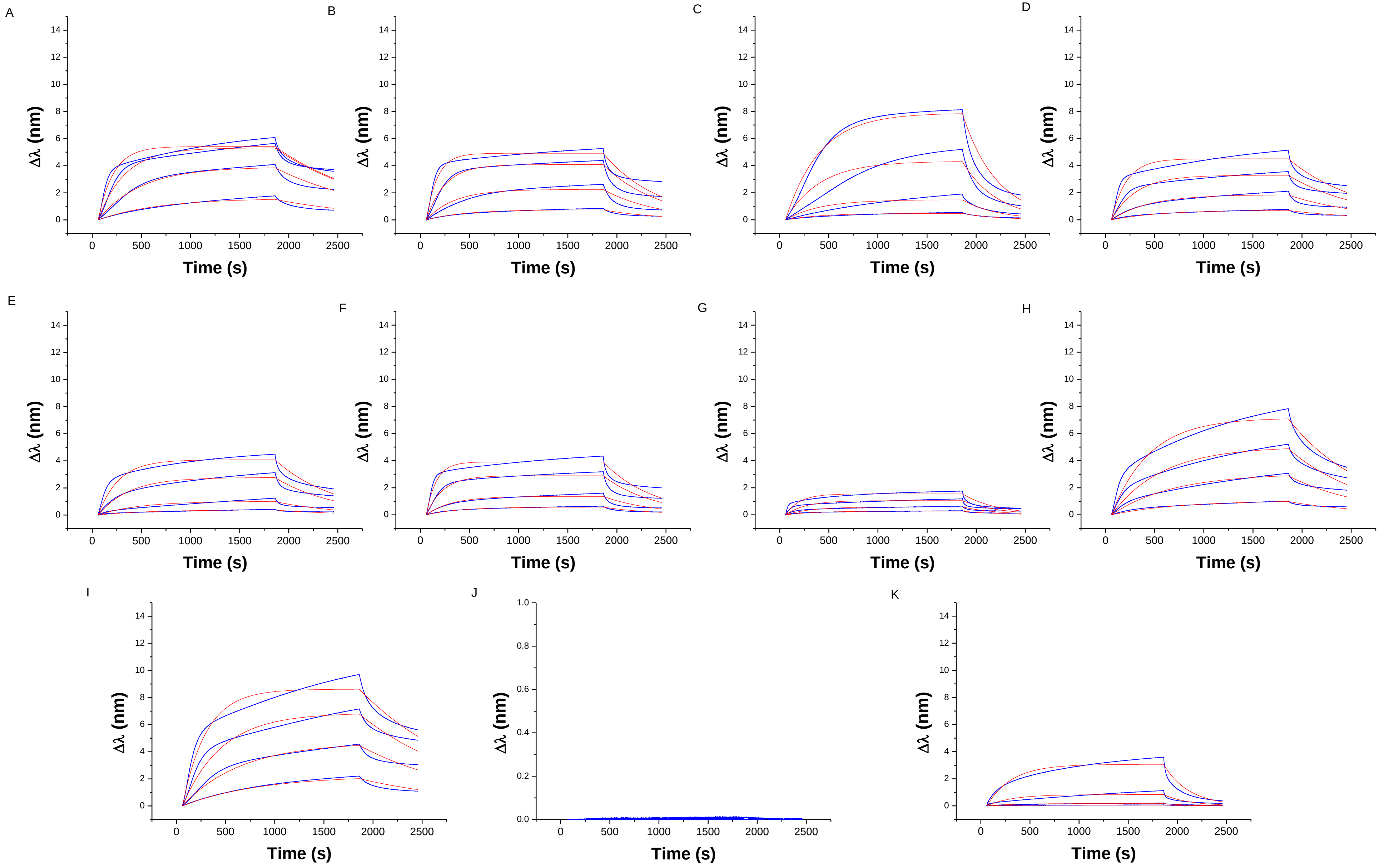


**Figure S5.** BLI analysis of binding of different concentrations of iMab scFv-$His_6$-FLAG (62.5, 125, 250 and 500 nM) to hTeloC 3x4 (**A**), hTeloC 3x3 (**B**), hTeloC X3 (**C**), hTeloC 2x3 (**D**), hTeloC 3x2 (**E**), hTeloC 2x3T (**F**), hTeloC 1x3T (**G**), hTeloC 1x6T (**H**), hTeloC 2x6T (**I**), hTeloC-Mut (**J**), and hTeloC-scr (**K**) at pH 7.5. Blue lines are experimental curves; red lines are the fits to the 1:1 model.

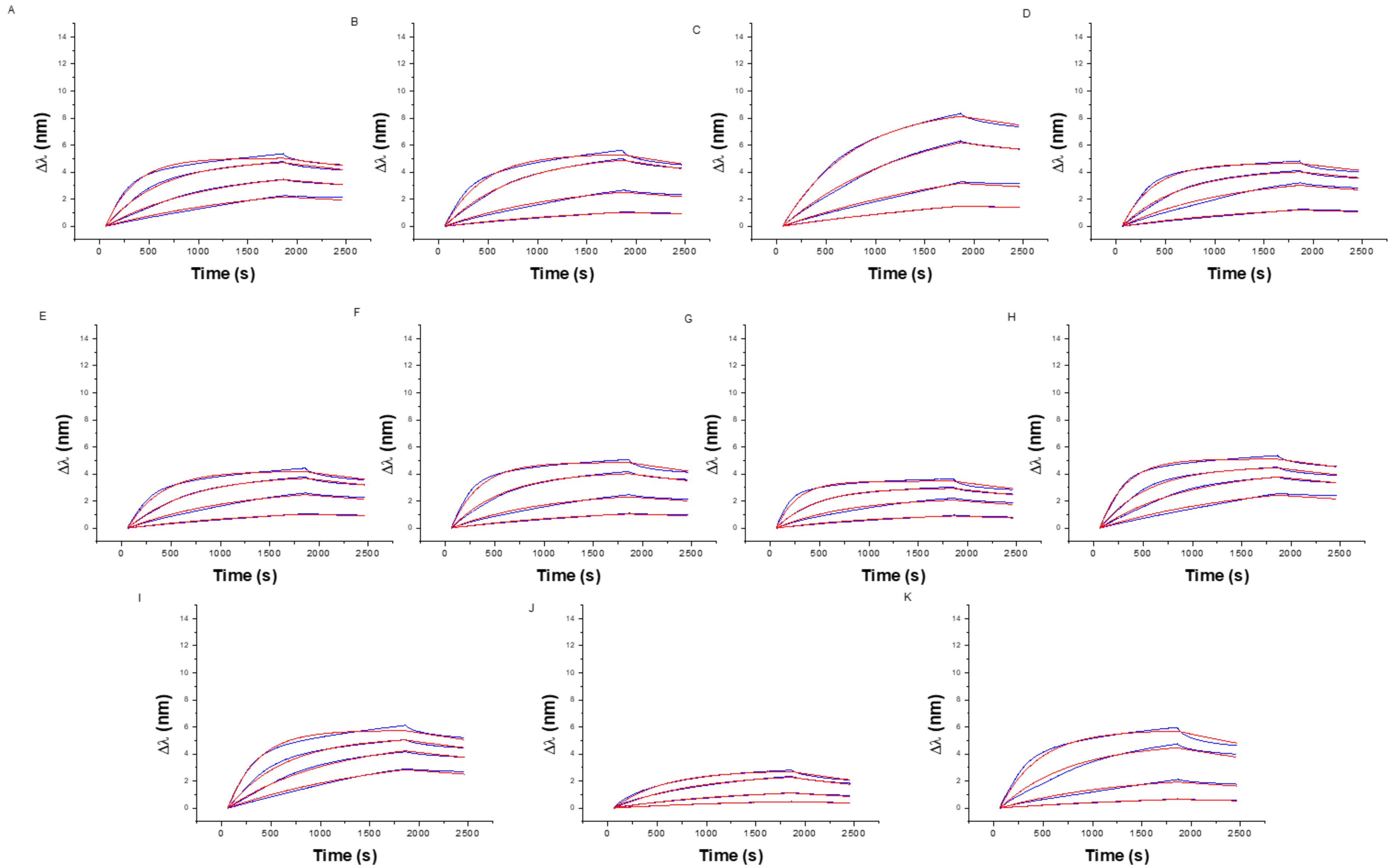


**Figure S6.** BLI analysis of binding of different concentrations of iMab scFv-His$_6$ (62.5, 125, 250 and 500 nM) to hTeloC (**A**), hTeloC 3x3 (**B**), hTeloC X3 (C**),** hTeloC 2x3 (**D**), hTeloC 3x2 (**E**), hTeloC 2x3T (**F**), hTeloC 1x3T (**G**), hTeloC 1x6T (**H**), hTeloC 2x6T (**I**), hTeloC-Mut (**J**), and hTeloC-scr (**K**) at pH 6.0. Blue lines are experimental curves; red lines are the fits to the 1:1 model.

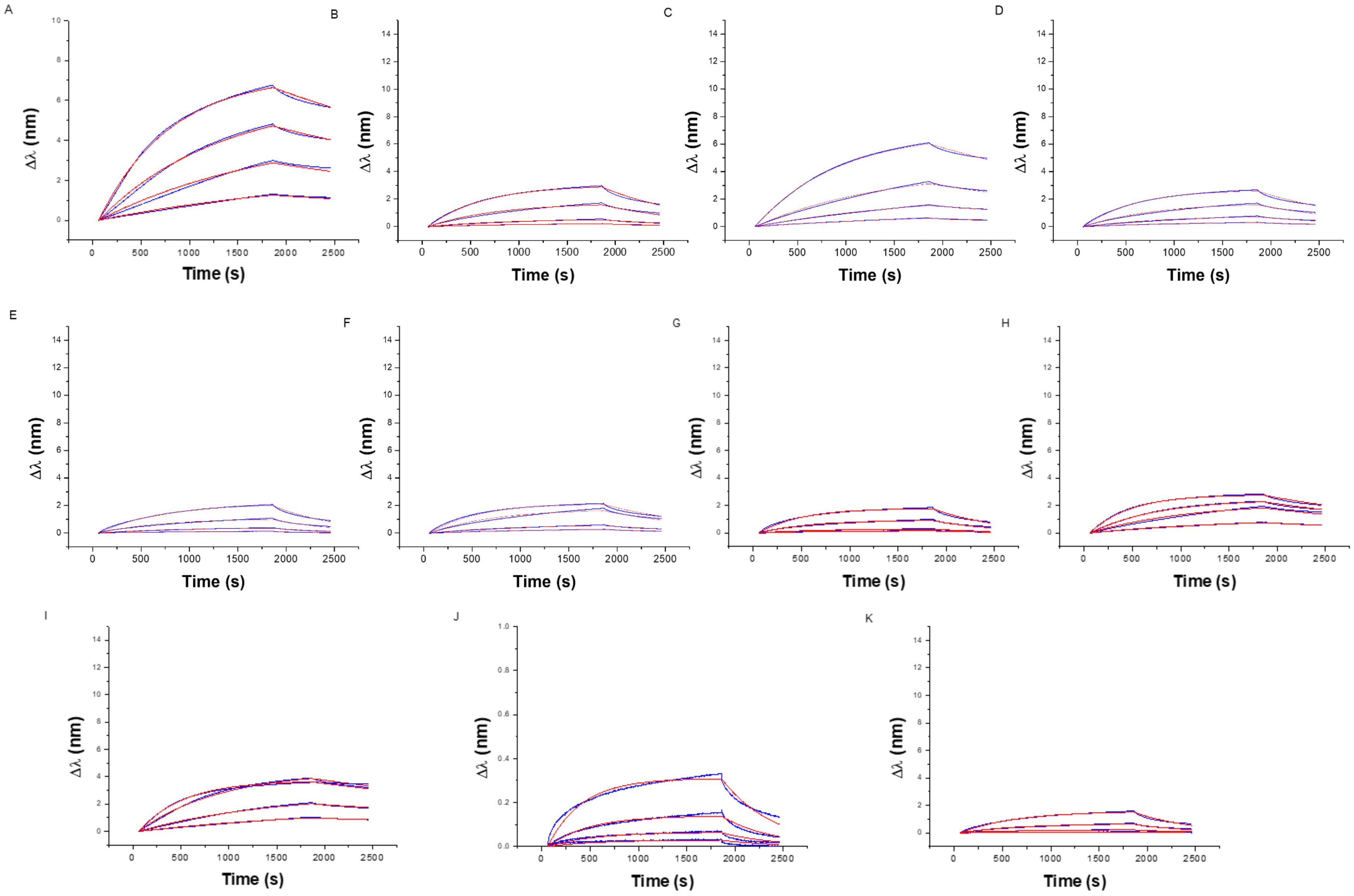


**Figure S7.** BLI analysis of binding of different concentrations of iMab scFv-His$_6$ (62.5, 125, 250 and 500 nM) to hTeloC 3x4 (**A**), hTeloC 3x3 (**B**), hTeloC X3 (**C**), hTeloC 2x3 (**D**), hTeloC 3x2 (**E**), hTeloC 2x3T (**F**), hTeloC 1x3T (**G**), hTeloC 1x6T (**H**), hTeloC 2x6T (**I**), hTeloC-Mut (**J**), and hTeloC-scr (**K**) at pH 7.5. Blue lines are experimental curves; red lines are the fits to the 1:1 model.

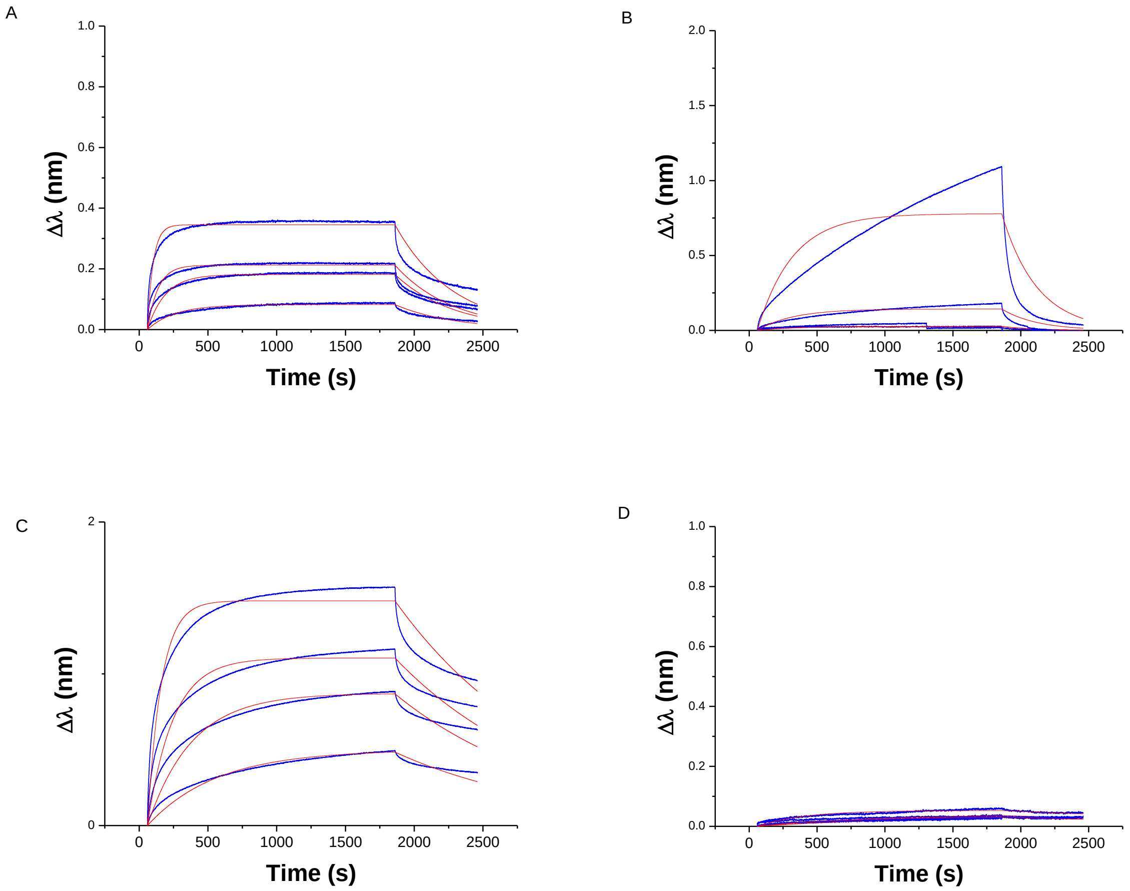


**Figure S8.** Bio-layer interferometry (BLI) analysis of binding of different concentrations of iMab scFv-His$_6$-FLAG at pH 6 (**A** and **C**) and 7.5 (**B** and **D**) for hp-ATT (**A**–**B**) and ss-DNA (**C**–**D**). The red lines are the fitting of the experiment. Concentration range: 62.5, 125, 250 and 500 nM.

#### Circular dichroism (CD) studies

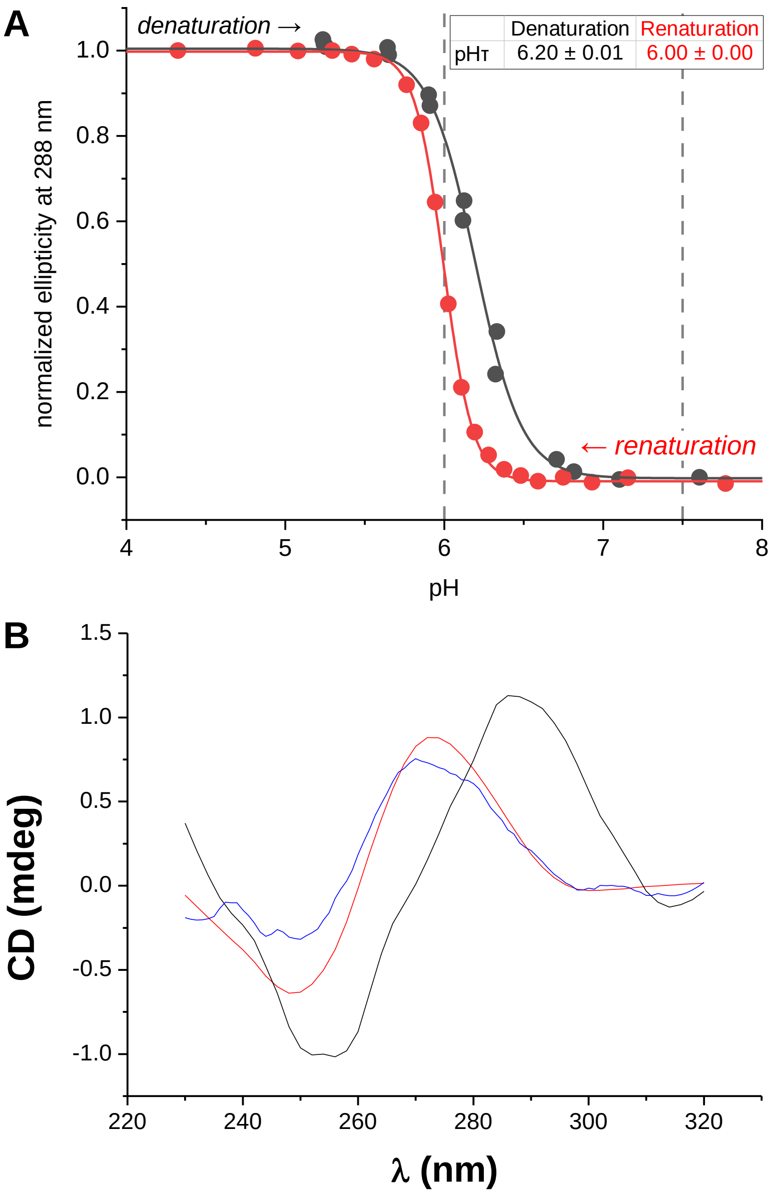


**Figure S9.** (**A**) pH-dependent denaturation/renaturation curves of hTeloC (2.5 µM in 10 mM $LiAsMe_2O_2$, 100 mM KCl buffer). Titrations were performed as described elsewhere (F. Berthiol et al., *Molecules* **2023**, *28*, 682). Dashed lines indicate the two conditions (pH 6.0 and 7.5) used in BLI experiments. (**B**) CD spectra of native hTeloC at pH 6.0 (black), pH 6.5 (blue) and pH 7.5 (red) at oligonucleotide concentration *c* = 2.5 µM.

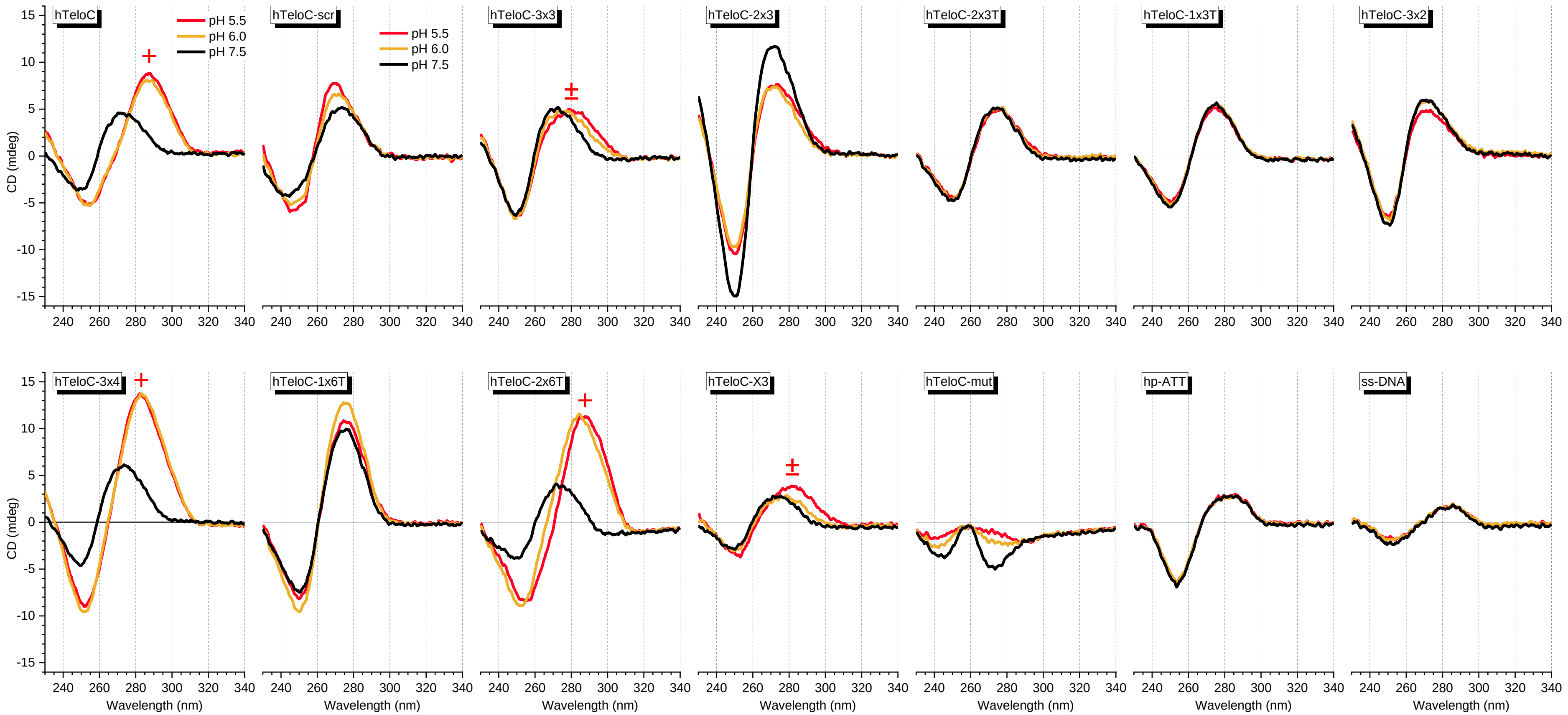


**Figure S10.** CD spectra of the sequences used in this study, obtained at pH 5.5 (red), 6.0 (dark yellow) and 7.5 (black). Oligonucleotide concentration $c$ = 2.5 μM in all cases. iM-characteristic peaks are labelled with a plus (+) symbol; peaks indicating partial formation of iMs are labelled with a plus-minus symbol (±). Note that the peculiar shape of hTeloC-mut spectrum is in agreement with literature data (M. Zeraati et al., *Nature Chem*. **2018**, *10*, 631).

### Bulk-FRET experiments

**Table S1.** Oligonucleotide sequences used in bulk-FRET experiments.

| Name | Sequence (5'→3') |
|---|---|
| Cy3-Py25 | **Cy3**-TTCCCCACCCTCCCCACCCTCCCCATGAGGACACGTGCATTCC |
| Cy3-hTelo | **Cy3**-TTAACCCTAACCCTAACCCTAACCCTAAATGAGGACACGTGCATTCC |
| Cy5-CS | GGAATGCACGTG**T***CCTC[a] |

T* = **Cy5**-modified dT residue

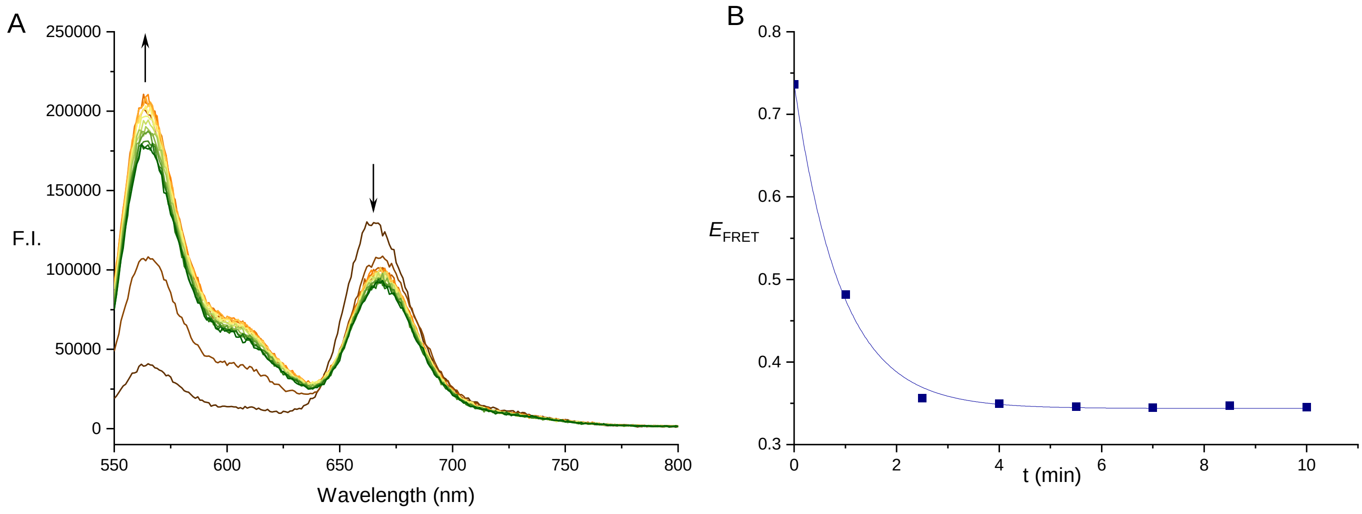


**Figure S11.** Time-dependent variation of fluorescence emission spectra (**A**) and $E_{FRET}$ value (**B**) of Cy3-Py25 / Cy5-CS substrate (4 nM) after addition of recombinant hnRNP K (800 nM) at pH 5.8. Spectra were recorded in 2-min intervals.
